# Preclinical immunogenicity of the LP.8.1-adapted BNT162b2 COVID-19 vaccine

**DOI:** 10.64898/2025.12.15.694478

**Authors:** Chaitanya Kurhade, Wei Chen, Weiqiang Li, Kristin R. Tompkins, Lyndsey T. Martinez, Swati Rajput, Emily Babiarz, Aaron Yam, Shin-Ae Lee, Shikha Shrivastava, Sarah O’Leary, Subrata Saha, Hui Yao, Li Hao, Todd Coffey, Carla Iris Cadima Couto, Alexander Muik, Raquel Munoz Moreno, Wesley Swanson, Pilar Mendoza Daroca, Uğur Şahin, Annaliesa S. Anderson, Kena A. Swanson, Pirada Suphaphiphat Allen, Kayvon Modjarrad

## Abstract

SARS-CoV-2 evolution toward antigenically distinct lineages drives escape from host immunity. JN.1 lineage derivatives have recently dominated the global epidemiologic landscape. In preclinical models, an LP.8.1-adapted BNT162b2 vaccine elicited higher serum neutralizing antibody responses against contemporary, circulating JN.1 sublineages, including the currently dominant XFG, as compared to JN.1-, KP.2- and XEC-adapted vaccines. These findings supported the selection of an LP.8.1-adapted vaccine for the composition of the 2025-26 COVID-19 vaccine formula.

## MAIN TEXT

COVID-19 continues to cause substantial disease burden, with illness, hospitalization and death rates rivaling or exceeding those of influenza ^1^. The evolution of SARS-CoV-2, the cause of COVID-19, toward escape from host immunity necessitates continuous evaluation of vaccine antigens to ensure that they closely match circulating virus lineages. Since 2022, licensed COVID-19 vaccines have been periodically adapted to elicit protective immunity against contemporary SARS-CoV-2 lineages and have demonstrated consistent clinical and public health benefit across diverse populations in real-world settings ^2-4^. Updates are intended to keep pace with major antigenic changes in the SARS-CoV-2 Spike (S) protein ^5-7^. The most recent antigenic shift occurred in late 2023, with the emergence of the BA.2.86 lineage and its JN.1 sublineage ^8^. This change prompted regulatory authorities to recommend COVID-19 vaccine updates to a JN.1 composition in some parts of the world and to the JN.1 sublineage, KP.2, in other regions. JN.1 sublineages have caused most cases of COVID-19 over the last two seasons and remain globally dominant ^9,10^. In that time, the virus has undergone extensive genetic and antigenic drift. Acquired mutations in the S protein have conferred improved viral fitness, as reflected by increased transmissibility and escape from neutralization by COVID-19 vaccine anti-sera ^11-13^.

The accelerated evolution of SARS-CoV-2 has primarily manifested in the insertion, deletion and substitution of amino acid residues in the receptor binding domain (RBD) and N-terminal domain (NTD) of the S protein. An interaction of the number and locations of residue changes in these domains generally has corresponded to the magnitude of immune escape. Two of the globally dominant JN.1 sublineages in early 2025 (**Supplementary Figure 1A**)—XEC and LP.8.1—differ from one another by eight residues in the S protein and by six and nine residues compared to JN.1, respectively (**Supplementary Figure 1B**). This exceeds the three-residue difference between JN.1 and KP.2 ^14^, the two recommended lineage compositions for 2024-2025 vaccine formula, but is far less than the 37 S residue changes between JN.1 and the prior season’s recommended vaccine variant, XBB.1.5. ^15^ The epidemiologic trends in the first half of 2025 and the observed trends in genetic drift prompted us to evaluate the immunogenicity of both XEC- and LP.8.1-adapted BNT162b2 vaccines, as compared to the JN.1- and KP.2-adapted BNT162b2 vaccines.

Neutralizing antibody responses elicited by LP.8.1- and XEC-adapted BNT162b2 vaccines were evaluated in female BALB/c mice in either a COVID-19 mRNA vaccine-experienced or vaccine-naïve background. Vaccine-experienced mice received two doses of the original (wild-type (WT) (Wuhan strain)) BNT162b2 vaccine as a primary series and one dose of the bivalent BNT162b2 (WT + Omicron BA.4/5) vaccine one month later. Mice were then vaccinated, one month after the third dose, with either JN.1-, KP.2-, XEC- or LP.8.1 -adapted BNT162b2 vaccines. Sera were collected prior to and after the final vaccination for assessment of neutralization of epidemiologically relevant JN.1 sublineage pseudoviruses (JN.1, KP.2, XEC, LP.8.1, LP.8.1.1, LF.7, NB.1.8.1, XFG) and the BA.3.2 saltation lineage (**Figure 1A**).

**Figure 1.**
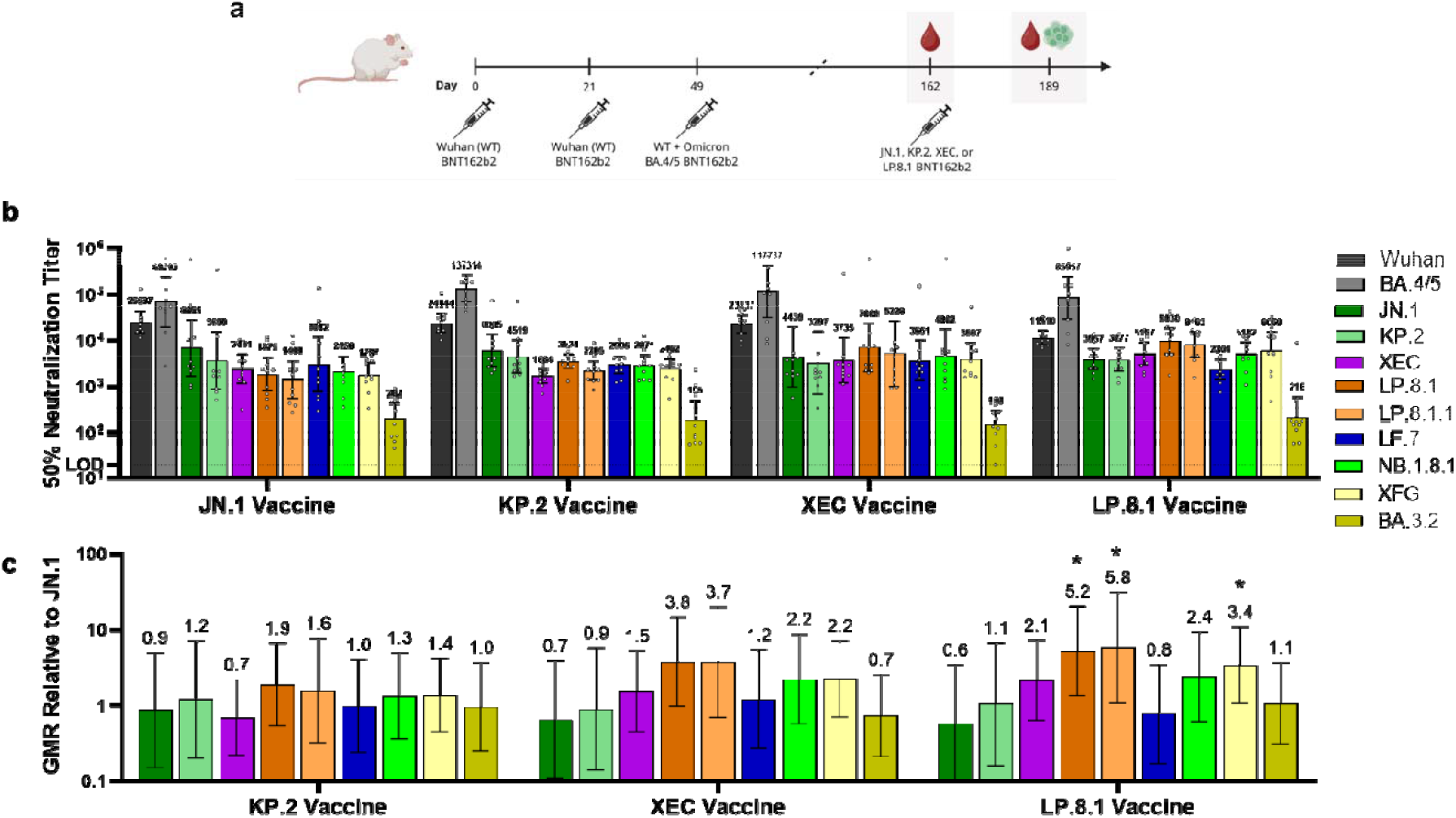
Pseudovirus neutralization titers elicited by JN.1-, KP.2-, XEC- and LP.8.1-adapted BNT162b2 vaccines administered to vaccine-experienced mice. **A**. Schematic for vaccination schedule including a fourth dose of the JN.1, KP.2, XEC and LP.8.1 BNT162b2 vaccine in vaccine-experienced mice. Female BALB/c mice (10/group) that were previously vaccinated with two doses of monovalent original WT BNT162b2, and one subsequent dose of bivalent WT + Omicron BA.4/5 received a single intramuscular dose of one of these vaccine regimens: JN.1, KP.2, XEC or LP.8.1. All vaccine formulations contained a total dose of 0.5_Jµg. Sera were collected before the fourth dose and one-month post-fourth dose. Spleens were harvested at one-month post-fourth dose from five mice. **B**. Fifty-percent geometric mean serum neutralizing titers were assessed in a pseudovirus neutralization assay one-month post-fourth dose against the WT reference strain Wuhan, the Omicron lineages and sublineages BA.4/5, JN.1, KP.2, XEC, LP.8.1, LP.8.1.1, LF.7, NB.1.8.1, XFG and BA.3.2. 50% pseudovirus neutralization titers are shown as geometric mean titers (GMT)_Jwith_J95% CI of 10 mice per vaccine group. Each data point represents one animal. **C**. The geometric mean ratio (GMR) is the GMT of individual pseudovirus responses of each vaccine group (JN.1, KP.2, XEC or LP.8.1) divided by the GMT of analogous pseudovirus responses of the JN.1 vaccine group. The limit of detection (LOD) is the lowest serum dilution, 1:20. Error bars represent 95% CIs. Asterisks indicate statistical significance of pseudovirus GMR relative to the corresponding pseudovirus in the JN.1 vaccine group as determined by ANOVA using a Dunnett’s multiple comparisons test. * p<0.05.

Prior to administration of the fourth dose, all lineage-adapted vaccines had similar neutralizing responses against vaccine-matched lineages (**Supplementary Figure 2A**). The highest neutralizing titers across the four vaccine groups were against the Wuhan and BA.4/5 lineages, reflecting the influence of immune imprinting, as observed in other COVID-19 vaccine-experienced animal studies ^16,17^. After the fourth dose, the LP.8.1-adapted vaccine elicited more than a two log-fold rise in geometric mean neutralization titers (GMTs) (geometric mean fold rise (GMFR)) against the matched LP.8.1 lineage. This was substantially higher than that observed for JN.1-, KP.2-, and XEC-adapted vaccines (**Supplementary Figure 2B**).

In mice given the JN.1 vaccine as a fourth dose, neutralizing responses against XEC and LP.8.1 were 2.8- and 3.7-fold lower than those elicited against the vaccine-matched JN.1 lineage (**Figure 1B**). By contrast, the LP.8.1-adapted vaccine elicited higher GMTs against most lineages tested, as compared to both JN.1- and KP.2-adapted vaccines (**Figure 1B**). The LP.8.1 vaccine generated 2.4-to-3.4-fold and 1.8-to-2.5-fold higher titers against currently subdominant NB.1.8.1 and dominant XFG lineages, as compared to JN.1 (**Figure 1C**) and KP.2 vaccines, respectively (**Supplementary Figure 3A**). Overall, neutralizing titers elicited by the LP.8.1 vaccine were significantly higher against LP.8.1 (p=0.02), its derivative LP.8.1.1 (p=0.04), and the currently dominant XFG lineage, compared to the JN.1 vaccine (p=0.04) (**Figure 1C**).S The XEC vaccine-elicited neutralizing titers were either similar or lower than those elicited by the LP.8.1 vaccine.

In contrast to a vaccine-experienced model, naïve mice typically generate more robust anamnestic responses to novel antigens, resulting in higher post-vaccination neutralizing titers. Responses elicited by naïve animals are also more useful for assessing antigenic distances between genetically distinct lineages. In this model, BALB/c mice received two doses of JN.1-, KP.2-, XEC- or LP.8.1-adapted BNT162b2 vaccines four weeks apart (**Figure 2A**). All vaccine groups exhibited robust neutralizing responses against the same pseudovirus panel used for vaccine-experienced mice (**Figure 1)**. As observed in prior studies ^16,17^, GMTs were approximately an order of magnitude greater than those observed in vaccine-experienced mice (**Figure 2B**). Neutralizing titers elicited by the LP.8.1-vaccine were 15.6-fold and 7.7-fold higher against the homologous LP.8.1, and 3.2-to-5.4-fold and 2-to-2.7-fold higher against other circulating JN.1 sublineages (XEC, LP.8.1.1, LF.7, NB.1.8.1, XFG), as compared to the JN.1 (**Figure 2C**) and KP.2 vaccines (**Supplementary Figure 3B**), respectively. The XEC vaccine, in contrast, elicited a smaller magnitude of relative increase in neutralizing responses at 1.9-to-3.3-fold times higher than the JN.1 vaccine (**Figure 2C**). Similar trends were observed relative to the KP.2 vaccine, wherein LP.8.1 and XEC vaccine-elicited neutralizing titers were respectively 2-to-7.7-fold and 0.9-to-2.2-fold higher (**Supplementary Figure 3B**). Overall, LP.8.1 vaccine titers were significantly higher against LP.8.1 (p<0.0001), LP.8.1.1 (p=0.003), LF.7 (p=0.008), NB.1.8.1 (p=0.018), and XFG (p=0.025) compared to those elicited by the JN.1 vaccine (**Figure 2C**). LP.8.1 vaccine-elicited neutralizing titers were also significantly higher against the homologous LP.8.1 sublineage (p<0.0001) and trended higher against other circulating JN.1 sublineages, relative to the KP.2 vaccine (**Supplementary Figure 3B**).

**Figure 2.**
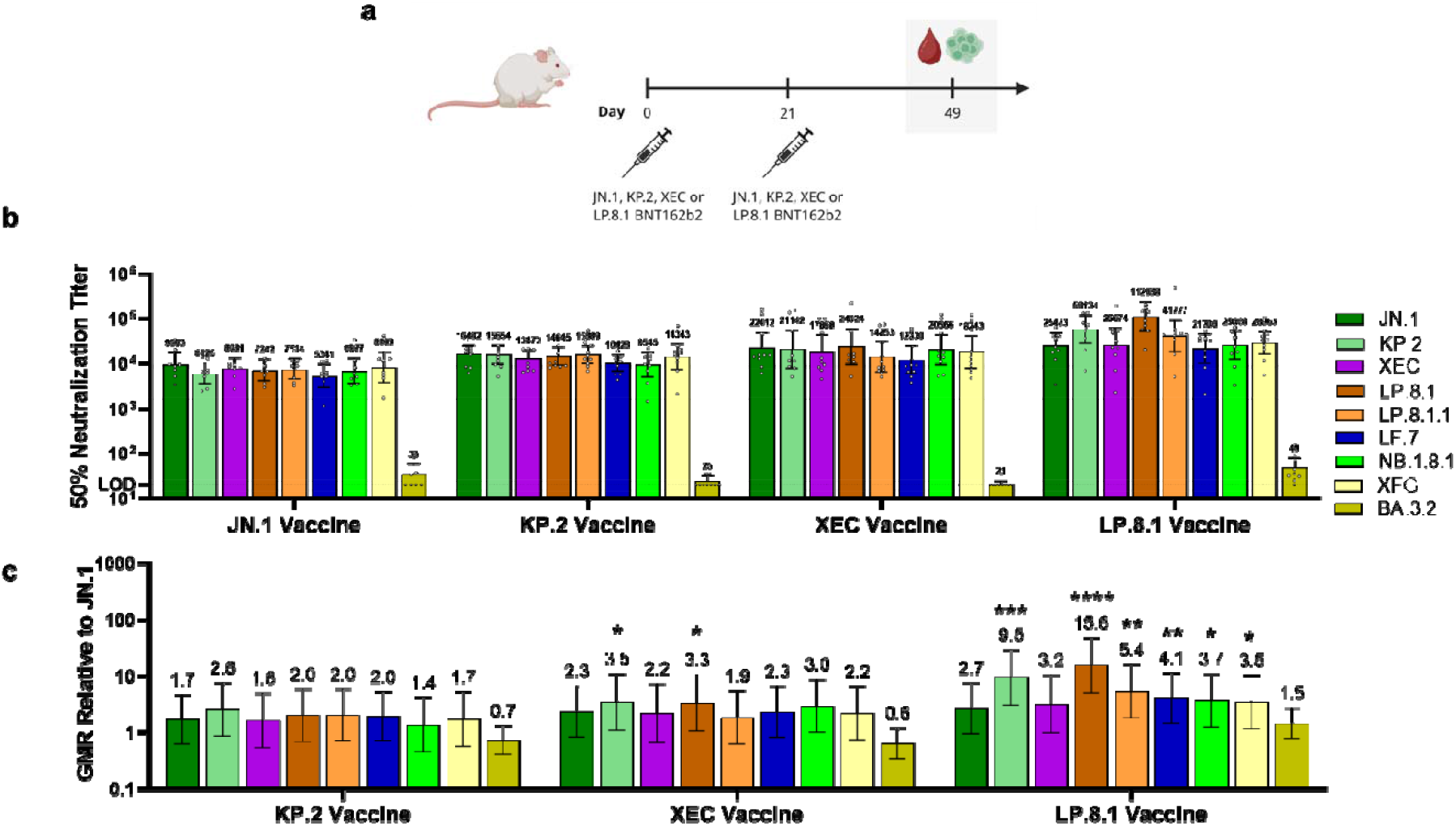
Pseudovirus neutralization titers elicited by JN.1-, KP.2-, XEC- and LP.8.1-adapted BNT162b2 vaccines administered as a primary series in naïve mice. **A**. Female BALB/c mice (10/group) were vaccinated with two doses of either JN.1-, KP.2, XEC- or LP.8.1-adapted vaccine, at a 21-day interval. All vaccine formulations contained a total dose of 0.5_Jµg. Sera and spleens were collected one-month after the second dose. **B**. Fifty-percent geometric mean serum neutralizing titers were assessed in a pseudovirus neutralization assay one-month after the second dose against JN.1, KP.2, XEC, LP.8.1, LP.8.1.1, LF.7, NB.1.8.1, XFG and BA.3.2 lineages and sublineages. 50% pseudovirus neutralization titers are shown as geometric mean titers (GMT)_Jwith_J95% CI of 10 mice per vaccine group. Each data point represents one animal. **C**. The geometric mean ratio (GMR) is the GMT of individual pseudovirus responses of each vaccine group (JN.1, KP.2, XEC or LP.8.1) divided by the GMT of analogous pseudovirus responses of the JN.1 vaccine group. The limit of detection (LOD) is the lowest serum dilution, 1:20. Error bars represent 95% CIs. Asterisks indicate statistical significance of pseudovirus GMR relative to the corresponding pseudovirus in the JN.1 vaccine group as determined by ANOVA using a Dunnett’s multiple comparisons test. ****p<0.0001, ***p<0.001, ** p<0.01, * p<0.05.

Although our assessment of breadth focused on dominant and subdominant JN.1 sublineages, the emergence of the saltation variant BA.3.2 in 2024 raised concerns as it harbors 44 amino acid differences in the S protein that are distinct from those of LP.8.1 and published reports have shown that these residue substitutions conferred neutralization escape from COVID-19 mRNA vaccine anti-sera ^11,18^. In our vaccine-experienced mouse model, we observed neutralizing titers against BA.3.2 that were 34-, 23-, 25-, and 46-fold lower than vaccine-matched lineages in the respective JN.1, KP.2, XEC and LP.8.1 vaccine groups (**Figure 1B**). In naïve mice, neutralizing titers against BA.3.2 were at or near the assay limit of detection (LOD), indicating complete immune escape from all four lineage-adapted vaccines (**Figure 2B**). These data suggest that the BA.3.2 lineage may be able to escape host immunity in the general population.

We employed antigenic cartography to map neutralizing titers, represented as antigenic distance, elicited by lineage-adapted vaccines in naïve mice. In this model, JN.1-sublineages were within two antigenic units of each other, corresponding to a 4-fold difference in GMT (**Supplementary Figure 4A**). This indicates an incremental, but progressive, antigenic drift of more recent JN.1 sublineages from their parental lineage. BA.3.2, however, is approximately 8 antigenic units away from the LP.8.1 lineage, likely representing a major shift in the antigenic composition of the SARS-CoV-2 S protein, as it has a similar distance from earlier Wuhan, BA.1, BA.4/5, XBB.1.5 and JN.1 lineages.

T cell-mediated immunity elicited by the lineage-adapted vaccines was also assessed using peptide pools covering the full-length S protein from Wuhan, BA.4/5, JN.1, KP.2, XEC and LP.8.1 lineages. In both vaccine-experienced and naive mouse models, the LP.8.1 vaccine elicited Th1-biased, cross-reactive T cells across all lineages tested. All lineage-adapted vaccines elicited comparable frequencies of cytokine-expressing CD4^+^ and CD8^+^ T cells (**Supplementary Figure 5 & 6**). These responses trend similarly to those previously induced by the original and earlier lineage-adapted BNT162b2 formulas, where T cell responses were consistent across lineage compositions, irrespective of antigenic distance ^16,17^.

The progressive antigenic drift of SARS-CoV-2 presents an ongoing challenge to maintaining vaccine-induced protective immunity. Regular updates to the lineage composition of vaccines have helped mitigate this challenge. Preclinical data herein build on reports that recent JN.1 sublineages have drifted far enough antigenically to evade neutralization by approved JN.1- or KP.2-adapted vaccines ^11,13^. The LP.8.1-adapted BNT162b2 vaccine, in mice with varying immune backgrounds, elicited higher neutralization titers against the homologous LP.8.1 sublineage, previously dominant XEC sublineage, and contemporary sublineages NB.1.8.1 and XFG, as compared to JN.1-, KP.2- and XEC-adapted vaccines. The emergence of BA.3.2 and demonstrated immune escape reaffirms the persistent threat of antigenic shift events. As of December 2025, there have been 106 cases of sequence-confirmed BA.3.2 (**Supplementary Figure 7**)^19^. Although BA.3.2 has a reported lower infectivity and slower epidemiologic growth rate than other circulating lineages ^11^, its recently rising incidence in Australia and Europe justifies its close monitoring.

Overall, these data demonstrate that the LP.8.1-adapted BNT162b2 vaccine maintains potent cross-neutralization of currently and recently predominant SARS-CoV-2 lineages, particularly as compared to JN.1- and s XEC-adapted vaccines, which have been recommended and approved as an alternative vaccine composition in some regions.^20^ The data presented herein are consistent with prior reports in demonstrating immunologic lineage-adapted vaccine formula over the course of a season’s antigenic drift.

## METHODS

### Animal ethics

All mouse experiments were performed at Pfizer, Inc. (Pearl River, NY, USA), which is accredited by the Association for Assessment and Accreditation of Laboratory Animal Care (AAALAC). All procedures performed on animals were in accordance with regulations and established guidelines and were reviewed and approved by an Institutional Animal Care and Use Committee (IACUC) or through an ethical review process. Animals were observed after all procedures and injection sites were monitored following each vaccination. Per IACUC guidelines, prior to euthanasia, animals were induced to a surgical plane of anesthesia via inhalation of ∼2-4% isoflurane. Exsanguination was then performed via cardiac puncture, with cervical dislocation as a confirmatory method.

### BNT162b2 mRNA XEC and LP.8.1 vaccine modification and formulation

The BNT162b2-adapted vaccines encode the S(P2) of JN.1 (GISAID EPI_ISL_18374006), KP.2 (GISAID EPI_ISL_18912216), XEC (GISAID EPI_ISL_19283891), and LP.8.1 (GISAID EPI_ISL_19467828) on the BNT162b2 RNA backbone. Purified nucleoside-modified RNA was formulated into lipid nanoparticles as previously described by Maier et al ^21^.

### Immunogenicity in BNT162b2-experienced mice

Female BALB/c mice (10 per group, age 6–8 weeks; Jackson Laboratory) were vaccinated and bled as shown in **Figure 1A**. In brief, mice were vaccinated intramuscularly with a two-dose series (days 0 and 21) of a 0.5□µg dose level of original BNT162b2 WT vaccine, followed by a third dose booster (day 49) of bivalent WT + Omicron BA.4/5 vaccine, and a fourth dose booster (day 162) of either JN.1-, KP.2-XEC- or LP.8.1-adapted vaccines. Formulations contained a total dose of 0.5□µg. A control group of ten mice received saline injections in place of active vaccines. A total volume of 50□µL of vaccine or saline was administered intramuscularly to the upper outer hind leg for each animal. Sera were collected for evaluation of pseudovirus neutralizing antibody responses prior to the fourth dose (day 162) and at one-month post-vaccination timepoint (day 189). Spleens were collected on day 189 to evaluate cell-mediated immune responses.

### Immunogenicity in naïve mice

Female BALB/c mice (10 per group, age 6-8 weeks; Jackson Laboratory) were vaccinated and bled as shown in **Figure 2A**. Mice were vaccinated intramuscularly on days 0 and 21 with either JN.1-, KP.2-, XEC- or LP.8.1 -adapted vaccines. The control group received saline injections in place of active vaccine groups. Sera and spleens were collected 28 days after the second dose (day 49) for evaluation of pseudovirus neutralizing antibody responses and cell-mediated immune responses, respectively.

### Pseudovirus neutralization assay

Pseudovirus stocks were generated in HEK-293T cells (ATCC, ref.# CRL-3216) using SARS-CoV-2 spike plasmid DNA and vesicular stomatitis virus (VSV; VSVΔG(G)-GFP virus: Kerafast, ref.# EH1019-PM). VSV-based pseudoviruses used in the assay expressed the S protein from the following SARS-CoV-2 variants: WT (Wuhan-Hu-1, ancestral strain), BA.4/5, JN.1, KP.2, XEC, LP.8.1, LF.7, LF.7.2.1, NB.1.8.1 and XFG. Mouse immune sera were serially diluted and incubated with individual VSV-based pseudoviruses before infecting Vero cell monolayers. Sera antibody neutralization was detected and enumerated by fluorescent foci using a CTL Immunospot Analyzer (Cellular Technology Limited) and quantified as 50% neutralization titer (NT_50_). Assay details can be found in the Supplementary Materials.

### JN.1, KP.2, XEC and LP.8.1 Antigenic Cartography

Antigenic maps were constructed using the antigenic cartography toolkit Racmacs v1.2.9 (https://acorg.github.io/Racmacs/index.html) as previously described ^17^. Details on the cartography process and workflow can be found in the Supplementary Materials.

### Global prevalence analysis of JN.1 sublineages and BA.3.2

SARS-CoV-2 virus sequences were downloaded from GISAID ^19^ in November 2025 and XEC, LP.8.1, LF.7, NB.1.8.1, XGF, and BA.3.2 lineages were assigned using Nextclade ^22^. Each major lineage group includes this specific lineage name as labeled and its sublineages. Isolates were filtered by collection data starting from October 2024 or January 2025. Global prevalence trends were plotted using R.

### T-cell response assay

Spleens were harvested from vaccinated mice, processed into a single cell suspension, and stimulated *ex vivo* with overlapping peptides representing the full S protein from Wuhan (WT), BA.4/5 and JN.1 lineages and KP.2, XEC and LP.8.1 sublineages, then analyzed for activation markers and cytokine production in CD4^+^ and CD8^+^ T cells by flow cytometry. Data acquisition and quality control were performed on a Cytek Aurora system, with subsequent analysis using OMIQ® software. The gating strategy is shown in **Supplementary Figure 8**. Assay details can be found in the Supplementary Materials.

### Statistical Analysis

Immunogenicity titers from mouse sera were log_10_-transformed for analysis. Vaccine group means at the last post-vaccination timepoint were compared using pairwise contrasts resulting from a one-way ANOVA. Dunnett’s procedure was used on the contrasts to control the familywise significance level at 0.05 within each pseudovirus, using either the JN.1 or KP.2 vaccine as reference. Means and mean differences (and their 95% confidence intervals) on the log_10_ scale were inverse-transformed and reported as GMT and GMR, respectively. All analyses were performed with SAS v9.4 software.

## Data Availability

The primary data generated or analyzed during this study that support the findings are included in the main article and supplementary information file. Additional data are available from the corresponding author upon reasonable request and with permission from Pfizer Inc.

## Author contributions

C.K. wrote the original draft. K.M. and P.S.A. contributed to writing of the original draft, data interpretation, investigation and supervision of the work. K.M. provided conceptualization of the work and designed all studies. C.K., L.T.M., S.R. and W.L. contributed to data visualization.

C.K., W.C., W.L., K.R.T., L.T.M., S.R., E.B., A. Y., S.L., S. Shrivastava, S.O., S. Saha, H. Y., L. H., R.M.M., T.C. and P.M.D. contributed to investigation, methodology, data generation and interpretation. W.S., K.A.S., A.S.A. and U.S. contributed to supervising work. All authors contributed to review and development of the manuscript and have read and agreed to the published version.

## Acknowledgments

This study was supported by Pfizer. We thank Pfizer Vaccine Research and Development and Comparative Medicine colleagues and veterinary staff at Pearl River, NY for their contributions to the *in vivo* studies, assay development and implementation, material production and characterization, and material and technical support in molecular biology, virology, and immunology. The authors thank Teresa Haugel for reviewing the manuscript, Christina D’Arco for scientific writing, editing, and graphical support, and Charlotte Ford for editorial assistance.

