## Supplementary Material for "Preclinical immunogenicity of the LP.8.1-adapted BNT162b2 COVID-19 vaccine"

Chaitanya Kurhade et al.

### Supplementary Methods

#### Pseudovirus neutralization assay

Serial dilutions of heat-inactivated murine sera (3-fold) were incubated with pseudovirus (VSVΔG(G)-GFP expressing SARS-CoV-2 S protein) for 1 h at 37 °C before inoculating confluent Vero (ATCC, ref.# CCL81.2) cell monolayers in 96-well plates. Fluorescent virus-infected foci were detected 19–21 h after inoculation with an anti–VSV polyclonal antibody (Imanis Life Sciences, ref# REA005) and Alexa488-conjugated secondary antibody (Invitrogen, ref# A-11008) and enumerated using a CTL Immunospot Analyzer (Cellular Technology Limited). A 50% neutralization titer ($NT_{50}$) was calculated as the last reciprocal serum dilution at which 50% of the virus is neutralized compared to wells containing virus only. Each serum sample dilution was tested in duplicate. The assay titer range was 20 to 43,740. Any serum sample that yielded a titer >43,470 was prediluted and repeated to extend the upper titer limit; sera that failed to neutralize at the lowest serum dilution (1:20) were reported to have a neutralizing titer of 20 (lower limit of detection, LLOD). VSV-based pseudoviruses used in the assay expressed the S protein from the following SARS-CoV-2 lineages: Wild type (WT) (Wuhan-Hu-1, ancestral strain), BA.4/5, JN.1, KP.2, XEC, LP.8.1, LF.7, LF.7.2.1, NB.1.8.1, XFG and BA.3.2.

#### JN.1, KP.2, XEC and LP.8.1 antigenic cartography

Antigenic cartography is a method to quantify and visualize viral neutralization data. An antigenic map takes titration data that measures strength of reactivity of a group of antisera (sera where an individual mouse has been vaccinated with a unique antigen of a particular SARS-CoV-2 lineage) against a group of antigens (different SARS-CoV-2 lineages). Antigenic mapping uses multidimensional scaling to position antigens (viruses) and sera in a map to represent their antigenic relationships. Antigenically similar strains are spatially close to one another on the map, while antigenically distinct strains are further apart. The spacing between grid lines is one unit of antigenic distance, corresponding to a two-fold dilution of antiserum in the neutralization assay. The maps were constructed by Racmacs package in R using 2000 optimizations, with the minimum column basis parameter set to “none.”

#### T-cell response assay

Murine splenocytes were stimulated *ex vivo* with DMSO only (unstimulated) or specific amino acid (aa) peptide libraries (15aa, 11aa overlap, 1 to 2 µg/mL/peptide) representing SARS-CoV-2 S amino acid sequences. Six individual peptide pools represented the full-length S sequence of the ancestral Wuhan (WT) (JPT), BA.4/5 (JPT), JN.1 (JPT), KP.2 (Mimotopes), XEC (Mimotopes) and LP.8.1 (Mimotopes) lineages. Following stimulation, splenocytes were stained for CD154- (CD40L), IFN-γ-, TNF-α-, IL-2- and IL4-positive $CD4^+$ and $CD8^+$ T cells, as previously described [1]. Samples were acquired on a 5-Laser Aurora system (Cytek ®) using SpectroFlo® software (version 3.1.2). The instrument was subject to daily quality control procedures using SpectroFlo® QC Beads per manufacturer recommendations. Acquired data files were analyzed using OMIQ® software.

**a**

SARS-CoV-2 S amino acid position

NTD: 22–190; RBD: 346–493

| | 22 | 31 | 59 | 182 | 184 | 186 | 190 | 346 | 435 | 444 | 445 | 456 | 475 | 478 | 487 | 493 | 572 | 679 | 1086 | 1104 |
|---|---|---|---|---|---|---|---|---|---|---|---|---|---|---|---|---|---|---|---|---|
| JN.1 | T | S | F | K | G | F | R | R | A | K | H | F | A | K | N | Q | T | K | K | V |
| KP.2 | T | S | F | K | G | F | R | T | A | R | H | L | A | K | N | Q | T | K | K | L |
| XEC | N | S | S | K | G | F | R | R | A | R | H | L | A | K | N | E | T | K | K | L |
| LP.8.1 | T | | F | K | G | L | S | T | A | R | R | L | A | K | N | E | T | K | R | L |
| LP.8.1.1 | T | | F | K | G | L | S | T | A | K | R | L | A | K | N | E | T | R | R | L |
| LF.7 | N | P | F | R | G | F | S | T | A | R | H | L | A | K | N | Q | T | K | K | V |
| NB.1.8.1 | N | S | S | K | S | F | R | R | S | K | H | L | A | I | N | E | T | K | K | V |
| XFG | N | P | F | R | G | F | S | T | A | R | R | L | A | K | D | E | I | K | K | V |

**b**

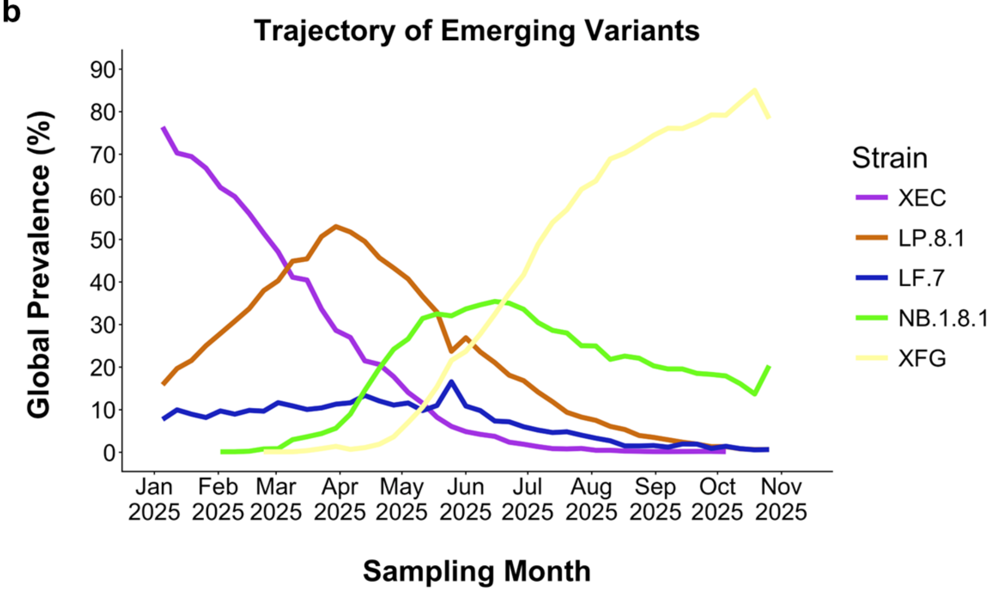

**Supplementary Figure 1. SARS-CoV-2 lineage Spike amino acid alignment and epidemiology**. **A.** Amino acid residues in the S protein are shown for the SARS-CoV-2 JN.1 lineage and relevant JN.1-derived sublineages (KP.2, XEC, LP.8.1, LP.8.1.1, LF.7, NB.1.8.1, and XFG). Blank squares represent deletions, and the darker blue squares indicate residue differences as compared to JN.1 lineage S protein. Amino acid positions belonging to the S N-terminal domain (NTD) and receptor binding domain (RBD) are indicated underneath their respective line. Created in BioRender. **B.** The trajectories of emerging JN.1-derived sublineages were plotted using SARS-CoV-2 sequences from the GISAID EpiCoV database [2] based on global prevalence (% of total global isolates in GISAID) from January through November 2025. Each colored line includes the labeled parental lineage (XEC, LP.8.1, LF.7, NB.1.8.1, and XFG) and its sublineages.

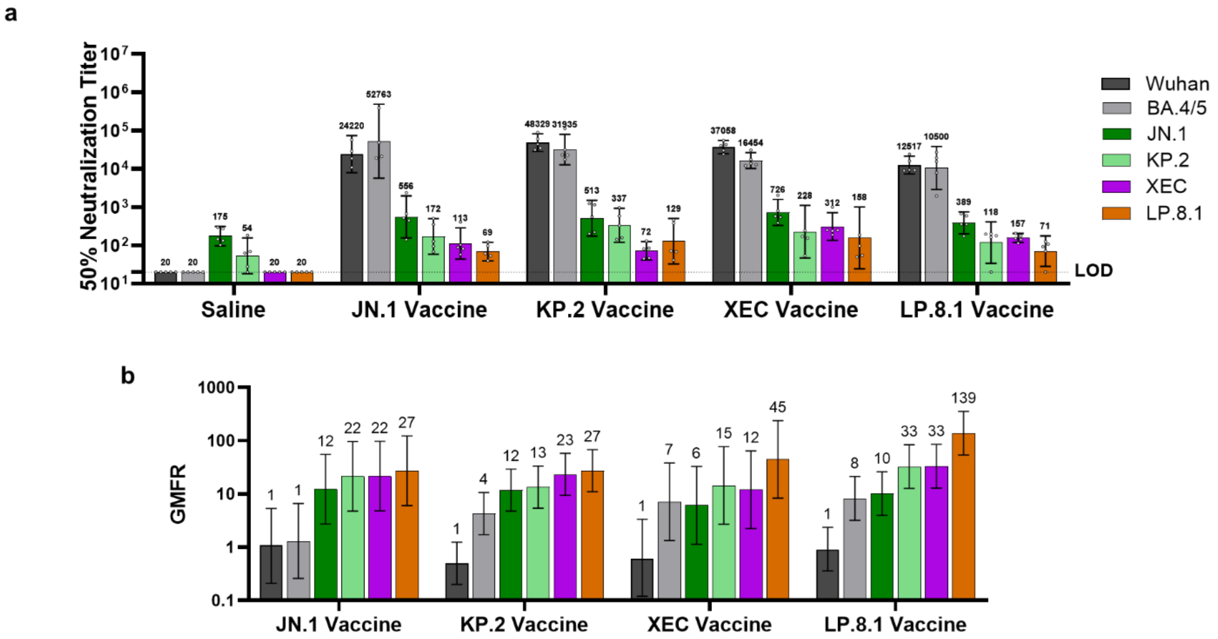

**Supplementary Figure 2. Pseudovirus neutralization titers pre- to post-fourth dose with a BNT162b2-lineage adapted vaccine in vaccine-experienced mice. A.** Baseline pseudovirus neutralization titers ($NT_{50}$) prior to BNT162b2 lineage-adapted vaccine booster immunization in BNT162b2- experienced mice. Female BALB/c mice (10/group) that were previously vaccinated with two doses of monovalent original WT BNT162b2, and one subsequent dose of bivalent WT + Omicron BA.4/5 received a fourth intramuscular dose of one of these vaccine regimens: JN.1, KP.2, XEC and LP.8.1. All vaccine formulations contained a total dose of 0.5 µg. Serum neutralizing antibody responses were assessed in a pseudovirus neutralization assay for the pre-4$^{th}$ dose and 1 month post-fourth dose timepoints against the WT reference strain Wuhan, and the BA.4/5, JN.1, KP.2, XEC and LP.8.1 lineages and sublineages. 50% pseudovirus neutralization titers are shown as geometric mean titers (GMT) with 95% CI of 10 mice per vaccine group. Each data point represents one animal. **B.** The fold rise in geometric mean neutralizing titers (GMFR) from pre-fourth dose to one-month post-fourth dose are shown with 95% CI. for the WT reference strain Wuhan, BA.4/5, JN.1, KP.2, XEC and LP.8.1. The limit of detection (LOD) is the lowest serum dilution, 1:20.

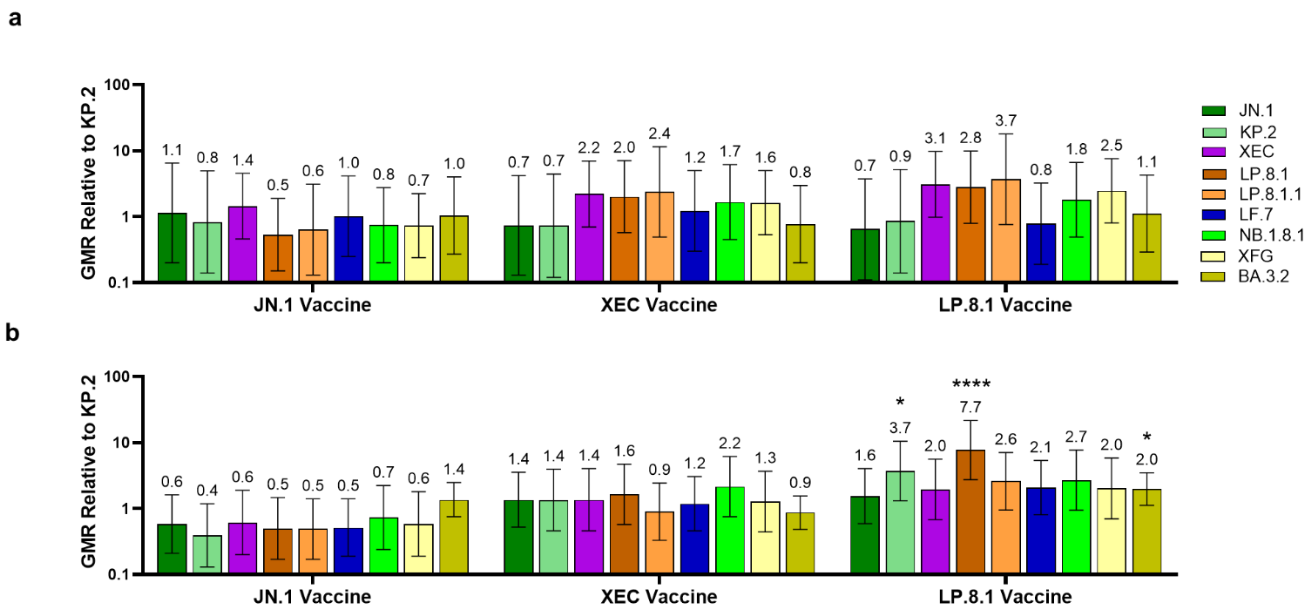

**Supplementary Figure 3. Neutralizing antibody response of vaccine-experienced and naïve vaccinated groups (JN.1, KP.2, XEC or LP.8.1) relative to KP.2 vaccine group. A.** Female BALB/c mice (10/group) that were previously vaccinated with two doses of monovalent original WT BNT162b2, and one subsequent dose of bivalent WT + Omicron BA.4/5 received a fourth intramuscular dose of one of these vaccine regimens: JN.1, KP.2, XEC and LP.8.1. Sera were collected one-month post-fourth dose. **B.** Female BALB/c mice (10/group) were vaccinated with two doses of one of the following vaccine regimens at a twenty-one-day interval: JN.1, KP.2, XEC and LP.8.1. Sera were collected one-month post-second dose. All vaccine formulations contained a total dose of 0.5 µg. Fifty-percent geometric mean serum neutralizing titers were assessed in a pseudovirus neutralization assay against the JN.1, KP.2, XEC, LP.8.1, LP.8.1.1, LF.7, NB.1.8.1, XFG and BA.3.2 lineages and sublineages. The geometric mean ratio (GMR) is the geometric mean titer (GMT) of individual pseudovirus responses of each vaccine group (JN.1, KP.2, XEC or LP.8.1), shown in Figure 1B and 2B, divided by the GMT of analogous pseudovirus responses of the KP.2 vaccine group. Error bars represent 95% CIs. Asterisks indicate statistical significance of pseudovirus GMR relative to the corresponding pseudovirus in the JN.1 vaccine group as determined by a two-sided ANOVA using a Dunnett's multiple comparisons test. ****$p<0.0001$, * $p<0.05$.

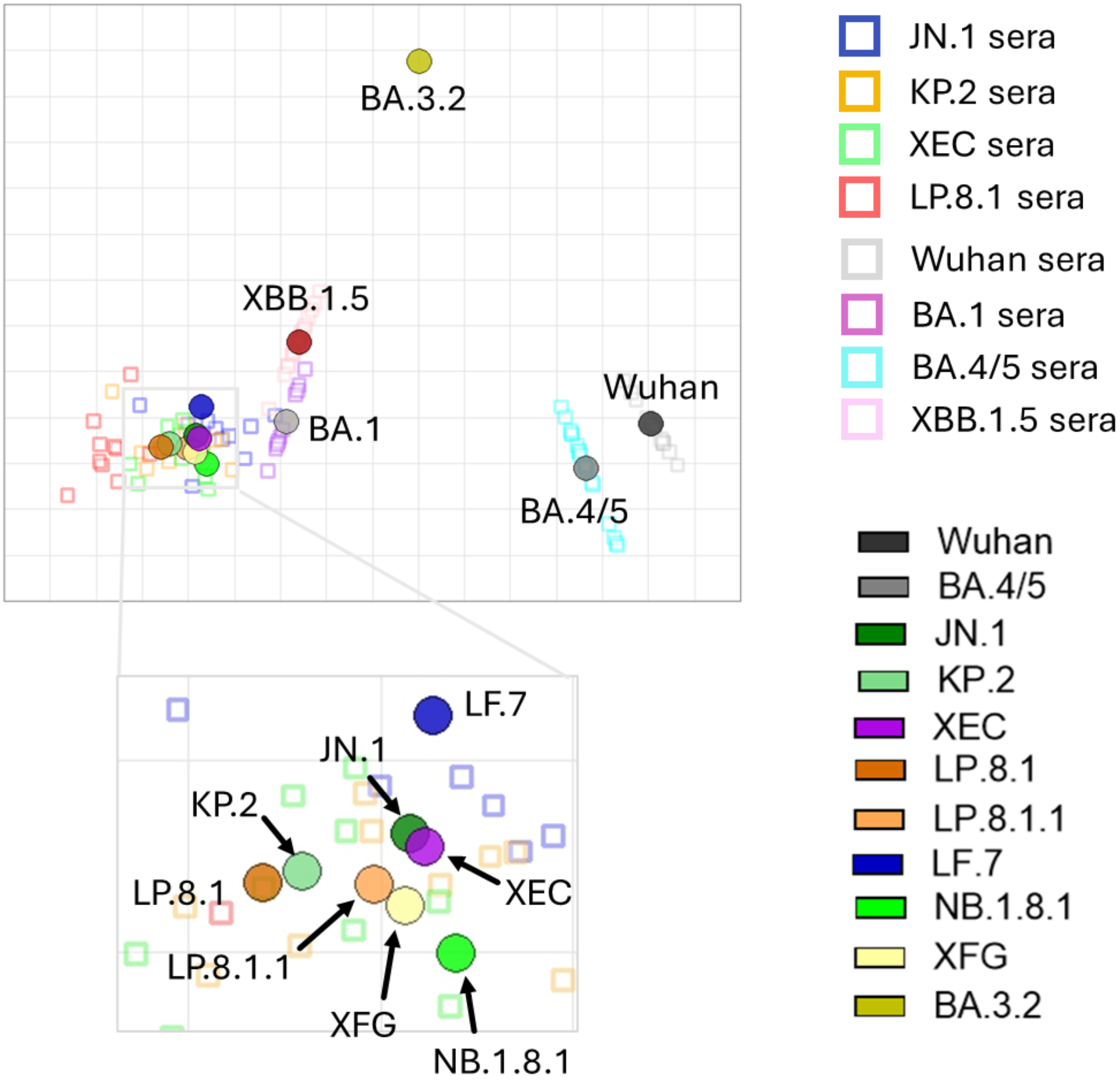

**Supplementary Figure 4. Antigenic map based on neutralization titers ($NT_{50}$) from sera of naïve mice immunized with JN.1, KP.2, XEC and LP.8.1 vaccines against indicated strains.** The antigenic map visualizes cross-reactivity among a panel of SARS-CoV-2 lineages using sera from vaccinated naïve mice. SARS-CoV-2 lineages are shown as circles and sera are indicated as squares. Each square corresponds to sera of one individual mouse and is colored by the vaccine that mouse received: BNT162b2 JN.1 (dark green), KP.2 (light green), XEC (purple) or LP.8.1 (orange). Antigenic distance is represented in both horizontal and vertical axes. Each square in the matrix represents 1 antigenic unit, which reflects a two-fold difference in neutralization titer. The points that are more closely together reflect higher cross-neutralization and are therefore antigenically more similar.

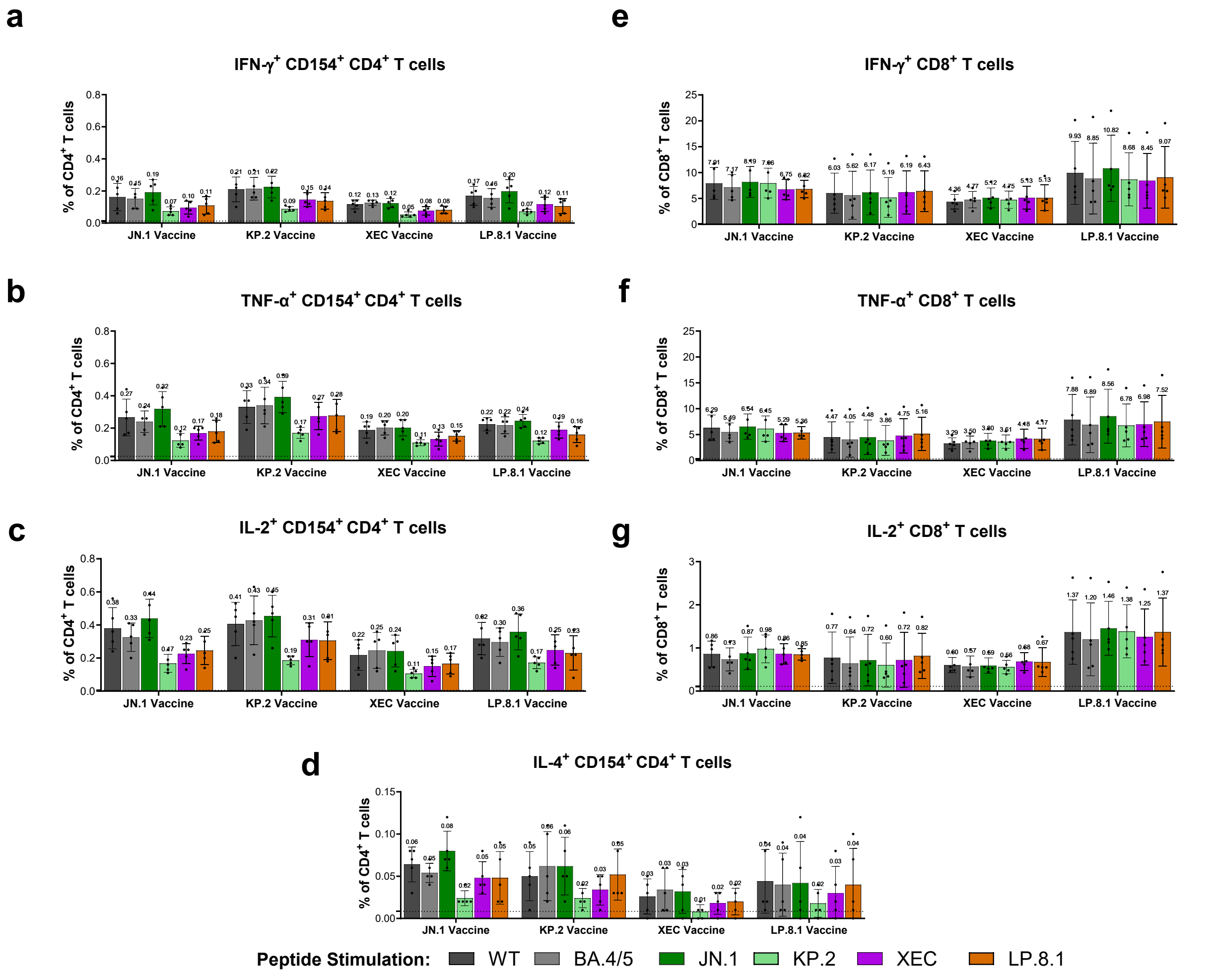

**Supplementary Figure 5. T cell responses elicited by BNT162b2 lineage-adapted vaccines administered as a fourth dose in BNT162b2-experienced mice.** One month following the fourth dose, splenocytes were harvested from mice (*n* = 5/group) and S-specific $CD4^+$ and $CD8^+$ T cells were characterized by a flow cytometry-based intracellular cytokine staining assay. All samples were stimulated separately with S peptide pools from the WT reference strain Wuhan and Omicron lineages and sublineages BA.4/5, JN.1, KP.2, XEC or LP.8.1. Graphs show the frequency of $CD4^+$ T cells expressing (**A**) IFN-γ, (**B**) TNF-α, (**C**) IL-2, and (**D**) IL-4, and the frequency of $CD8^+$ T cells expressing (**E**) IFN-γ, (**F**) TNF-α and (**G**) IL-2 in response to stimulation with each peptide pool across vaccine groups. Bars depict mean frequency ± SD.

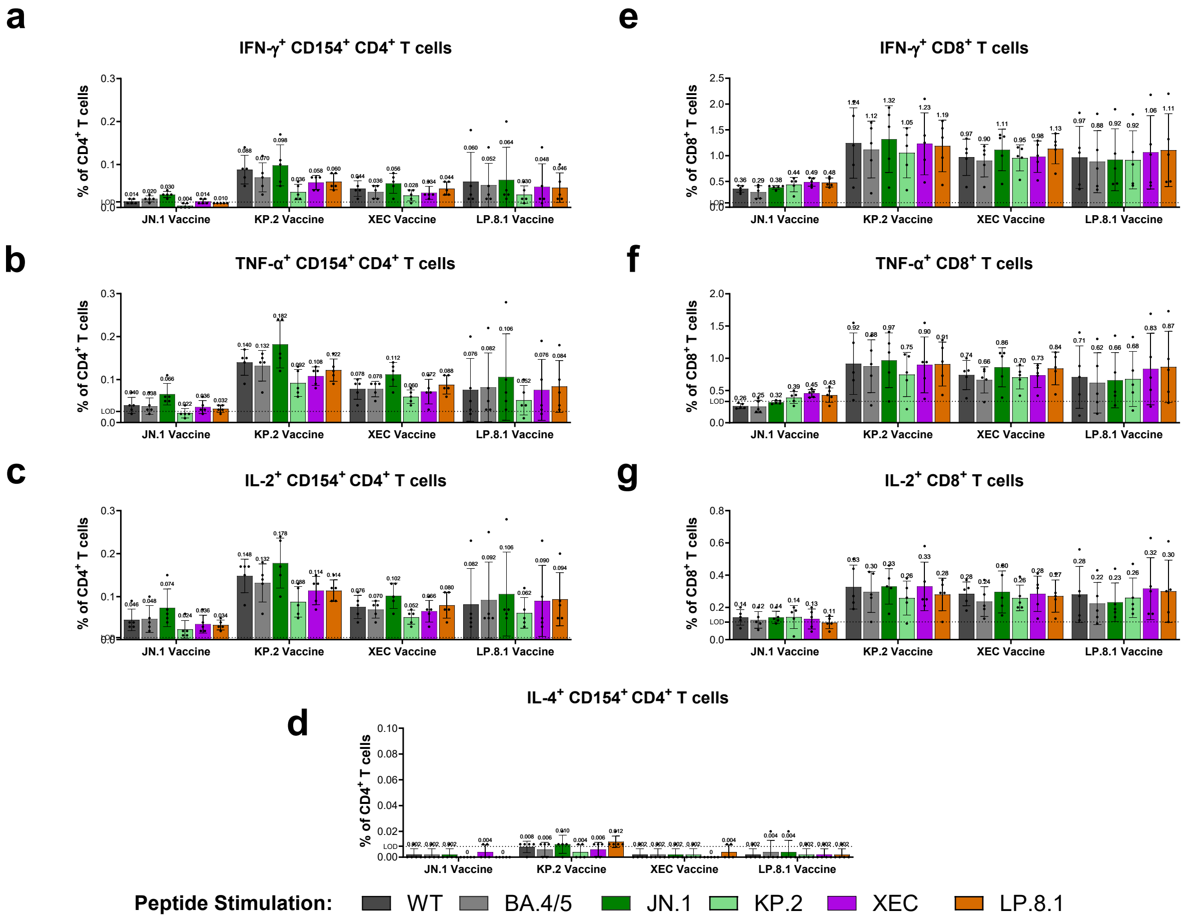

**Supplementary Figure 6. T cell responses elicited by BNT162b2 lineage-adapted vaccines administered as a primary series in naïve mice.** One month following the second dose (completion of primary series), splenocytes were harvested from mice (*n* = 5/group) and S-specific CD4$^+$ and CD8$^+$ T cells were characterized by a flow cytometry-based intracellular cytokine staining assay. All samples were stimulated separately with S peptide pools from the WT reference strain Wuhan and Omicron lineages and sublineages BA.4/5, JN.1, KP.2, XEC or LP.8.1. Graphs show the frequency of CD4$^+$ T cells expressing (**A**) IFN-γ, (**B**) TNF-α, (**C**) IL-2, and (**D**) IL-4, and the frequency of CD8$^+$ T cells expressing (**E**) IFN-γ, (**F**) TNF-α and (**G**) IL-2 in response to stimulation with each peptide pool across vaccine groups. Bars depict mean frequency ± SD.

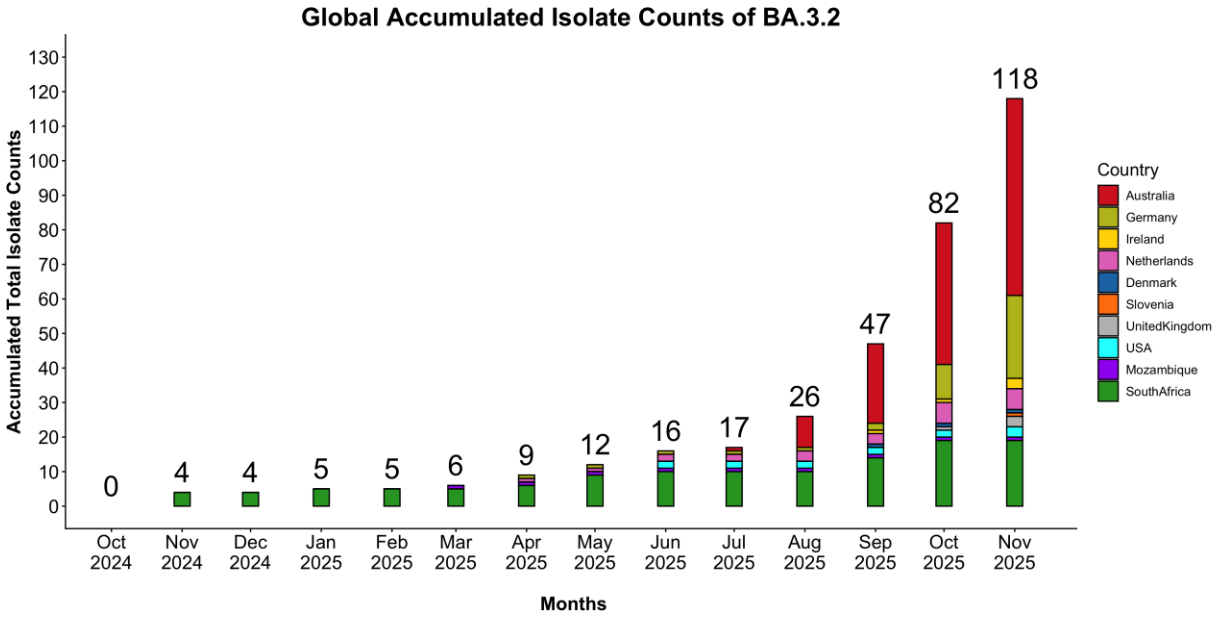

**Supplementary Figure 7. Global accumulated isolate counts of the SARS-CoV-2 BA.3.2 lineage by country**. SARS-CoV-2 virus sequences downloaded from GISAID [2] were filtered by collection and country data. Monthly isolate counts are shown for nine countries from October 2024 through November 2025.

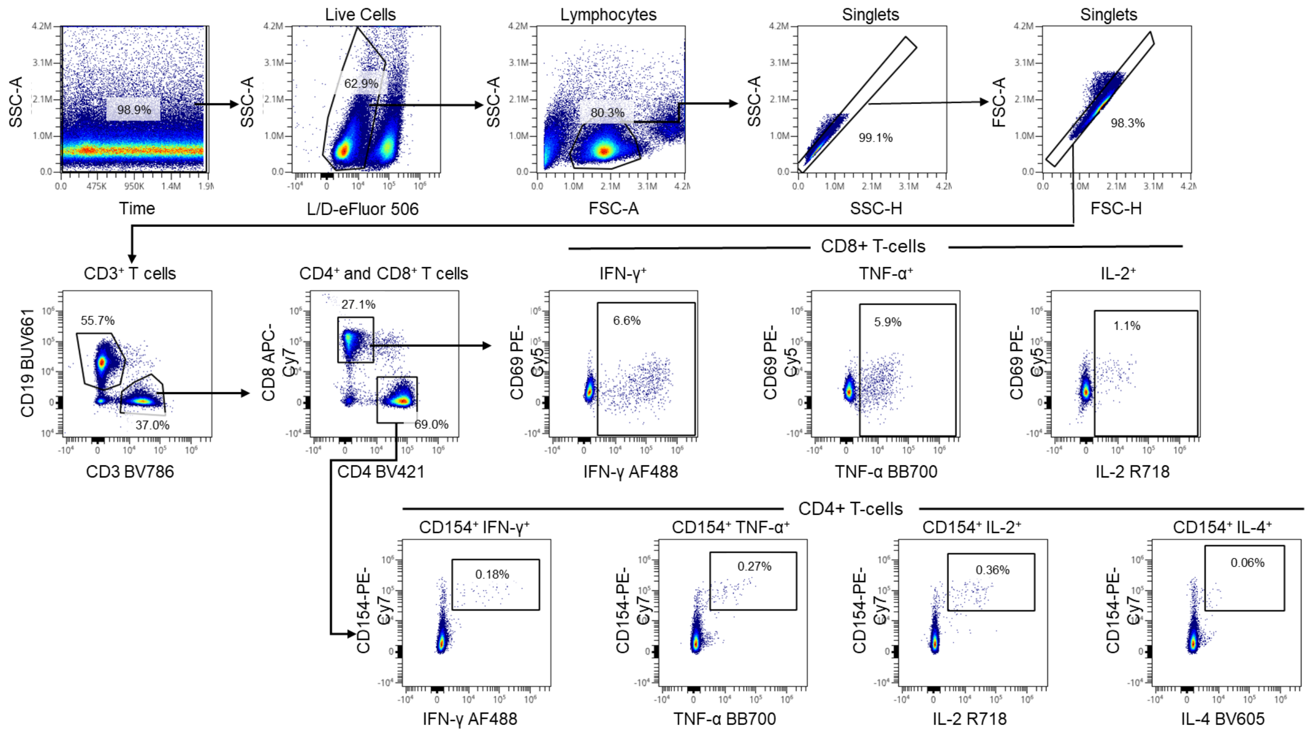

**Supplementary Figure 8. Gating strategy for intracellular cytokine staining flow cytometry analysis of T cell responses.** Flow cytometry gating strategy for identification of SARS-CoV-2 spike-specific T cells for different sublineages. (Upper row, left to right) Starting with events acquired with a constant flow stream and fluorescence intensity, viable cells, lymphocytes, and single events were identified and gated. Within singlet lymphocytes, $CD19^-$ $CD3^+$ T cells were identified and gated into $CD4^+$ and $CD8^+$ T cells (left middle row). Antigen-specific $CD4^+$ T cells were identified by gating on CD154 and cytokine-positive cells (bottom row). Activated $CD8^+$ T cells were identified by gating on CD69 and cytokine-positive cells (middle row). The antigen-specific cell frequencies were used for further analysis.
